# An electrogenetic toggle switch design

**DOI:** 10.1101/2022.05.19.492718

**Authors:** Lewis Grozinger, Elizabeth Heidrich, Ángel Goñi-Moreno

## Abstract

Synthetic biology uses molecular biology to implement genetic circuits that perform computations. These circuits can process inputs and deliver outputs according to predefined rules that are encoded, often entirely, into genetic parts. However, the field has recently begun to focus on using mechanisms beyond the realm of genetic parts for engineering biological circuits. We analyse the use of electrogenic processes for circuit design and present a model for a merged genetic and electrogenetic toggle switch. Computational simulations explore conditions under which bistability emerges in order to identify the circuit design principles for best switch performance. The results provide a basis for the rational design and implementation of hybrid devices that can be measured and controlled both genetically and electronically.

## 1 Introduction

Synthetic biology(*1–3*) engineers novel biological systems to fulfill predetermined functions using rational design, which depends fundamentally on mathematical models(*4–7*) and abstraction of the underlying biological processes. In particular, synthetic biology has developed sophisticated gene networks in bacteria (*8*) and other organisms(*9, 10*), which it has used for a variety of biotechnological applications from bioremediation(*11*) to biodiagnosis(*12*). The focus on genetic control is not accidental. Genetic networks regulate essential cellular process in bacteria, and the combination of experiments with synthetic genetic networks and mathematical modelling can yield critical insight into the biology of the cell (*13*).

Synthetic gene networks are often designed to process chemical inputs into chemical outputs according to some rules that implement a function of interest to the designer. We refer to this processing of inputs into outputs as a biocomputation(*14, 15*). Arriving at a network design that performs the desired function well is typically a hard problem, motivating the use of mathematical models, numerical analysis and simulations(*16*) to predict the performance characteristics of a specific network before time and resources are spent building it(*17, 18*).

Synthetic biology has focused primarily on optical methods of measuring the output of biological networks, especially during the testing phases of the design process. Typically the synthetic gene network is designed to express a fluorescent reporter protein whose activity can be measurement using equipment such as flow cytometers(*19*). In addition, synthetic biological networks have been engineered to utilise optical inputs, and the field of optogenetics takes advantage of light-sensitive characteristics of biological networks in order to regulate expression in synthetic gene networks(*20*). Light is a signal that can be both emitted and sensed easily with electronics, and the combination of optical inputs and outputs has allowed the development of hybrid electronic and biological systems for the closed-loop control of synthetic biological networks based on optogenetics(*21*).

However, bacteria are capable of using many different types of inputs and outputs in their natural biological networks. For example exoelectrogenic bacteria(*22*) are capable of sensing electronic inputs(*23*) in the form of electrochemical potentials, and of producing electronic outputs such as electrical current. These exoelectrogenic bacteria couple the oxidisation of a substrate to the reduction of a solid extracellular acceptor and find application in various bioelectrochemical systems, for example in generating electrical power or for evolving hydrogen (*24*). When these bacteria use an electrode as the electron acceptor, the resultant movement of charge from substrate to electrode can be detected as an output electrical current. The rate at which the bacteria metabolise substrate is therefore correlated with their current output. Synthetic gene networks have been engineered to exploit this relationship in exoelectrogens such as *Geobacter sulfurreducens*(*25*) and *Shewanella oneidensis*(*26*), by controlling expression of enzymes involved in key metabolic pathways. These networks offer the synthetic biologist genetic control of the bacteria’s electronic output.

Exoelectrogens respond to changes in electron acceptor potentials by using different metabolic and electron transfer pathways, and by regulating genetic expression (*27–29*). Controlling the electrical potential of an electrode can therefore provide exoelectrogens with an electrical input to which they can respond. For example, by coupling the electrical potential of an electrode to the activity of the redox-sensitive transcription factor SoxR in *Escherichia coli*, it has been demonstrated that electronic control of synthetic genetic networks can be achieved using exogenous redox mediators(*23*).

Since both genetic control of electronic output and electronic control of genetic input have been engineered separately, a logical next step for scaling up the complexity of electrogenic devices would be a synthetic biological network combining both mechanisms— this position underpins our current work. Such a network would take an electronic input and use synthetic genetic networks to process it into an electronic output. In this particular case, electronic input is provided by control of the electrical potential of the electrode used by the bacteria as an electron acceptor. The electronic output is the measurement of the electrical current produced by the bacteria. However, synthetic biology has only relatively recently begun to consider how electrogenic processes might be used to build novel biological networks(*30, 31*), and predictive computational models for the rational design of complex dynamical behaviours with electrogenic components are yet to be developed and tested.

Exoelectrogens such as *Geobacter* can colonise electrodes to form electroactive biofilms(*32*). An electroactive biofilm is composed of the bacteria themselves and an extracellular matrix with the capability of transporting electrons over large distances from bacteria deep in the biofilm to the electrode-biofilm interface. As a result, exogenous redox mediators are not required in order to connect bacteria with the electrode. Furthermore, biofilms can support larger populations of exoelectrogens by providing electrode access to bacteria without direct electrode contact. Nevertheless an electroactive biofilm has a finite capacity for charge (*33*) and cannot transport charge to the electrode at an arbitrary rate. Therefore it is possible that transport in the biofilm becomes the limiting step in current production and provides an upper bound on the potential depth of electroactive biofilms (*34*). Limiting transport in the biofilm also presents a design challenge for synthetic biologists, in that it leads to heterogeneity in the condition of different parts of the biofilm(*35*). For example, one region of the biofilm may be rich in electron donor substrate, while in another substrate is depleted entirely. This means the same synthetic biological network might be required to operate under different environmental conditions, adding complexity which further motivates the rational design of such networks using mathematical models.

Here we model the biofilm-electrode dynamics of a bioelectrochemical system where exoelectrogens form a biofilm on an electrode and consume substrate to produce current. The purpose of the model is to investigate the different dynamic behaviours that could be implemented by engineering the exoelectrogenic bacteria with synthetic gene networks, while using electrical signals as the input and output of the system. We will use bistability as a case study to demonstrate the usefulness of the model.

In a bistable system there are two stable steady states and the system rests in one of these two states indefinitely until induced by some external force to switch to the other. Bistability is a fundamental type of dynamics in both natural and synthetic biology, and in fact some of the earliest work in synthetic biology was toward engineering a bistable synthetic gene network called the ‘genetic toggle switch’ (36). Emergence of bistability in the genetic toggle switch, and generally in dynamic systems, requires certain conditions on the components of the switch; relatively few of the possible realisations of the genetic toggle switch will produce bistability. A major contribution of the original genetic toggle switch work was the development of a simple model which could be used to identify the conditions under which a novel gene network would exhibit bistability.

In the following we model a biofilm-electrode system that might be found in a bioelectrochemical system where exoelectrogens form a biofilm on an electrode and consume substrate from which they produce current. The purpose of the model is to aid in the design of an electrogenetic toggle switch, a bistable hybrid electrogenic-genetic system that could be implemented by engineering exoelectrogenic bacteria with synthetic gene networks. The system should be able to switch between two different levels of steady state electrical current output, using a transient change in electrode potential to induce switching. As in the study of the genetic toggle switch, we aim to use the model to predict conditions under which the synthetic gene network will exhibit bistability by using a mathematical model and steady state analysis.

## 2 Results

The diagram in Figure 1A shows the information flow throughout the electrogenetic toggle switch design. The inputs to the system are controlled by modifying the potential of the electrode (*V*), and the output is measured as the current (*I*) generated by the biofilm. In our model, bistability was achieved in the relationship between these two values (*I-V*)—in a similar way to bistability in the original genetic toggle switch(*36*), which was characterised by the level of the output (a fluorescent protein) as related to the inputs (chemical inducers). Inputs are processed into outputs by the interplay between a synthetic gene network (*a*) and charge transport (*q*). Specifically, this interplay was designed to be a negative feedback loop (Figure 1B); in what follows we describe the conditions under which the dynamics of the feedback loop facilitates the emergence of bistability.

**Figure 1:**
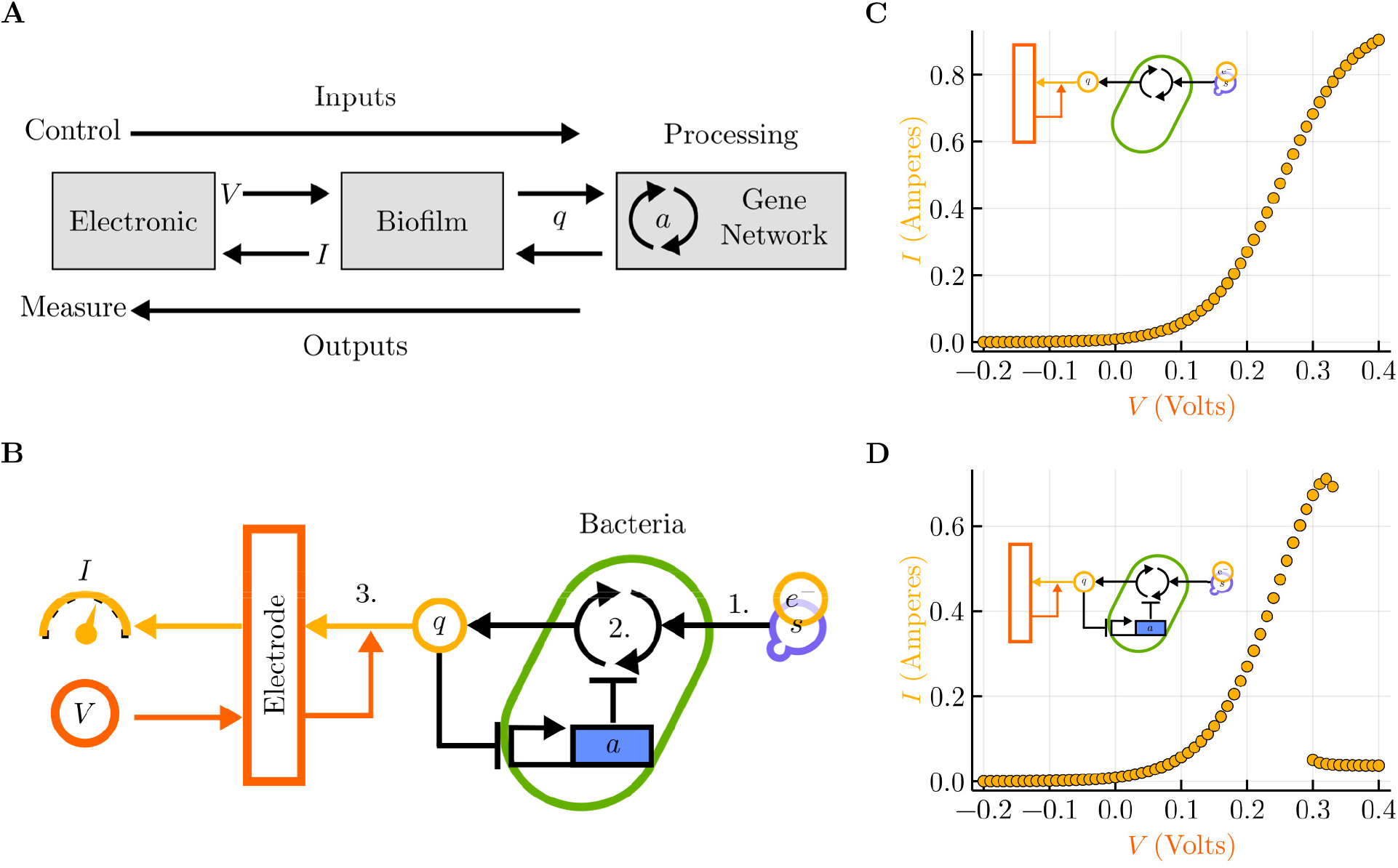
An overview of the electrogenetic toggle switch design. **A.** Diagram showing the key components of the system and information flow at a high level. Inputs to the system are provided by an electronic device controlling the potential (*V*) of an electrode which is transmitted by the biofilm to a gene network. The gene network processes the input to produce output as a variable level of charge by converting substrate *s* to charge *q* at a rate inhibited by a gene *a*. The output is the electronic current (*I*) generated by the biofilm. **B.** Detailed model of the negative feedback loop responsible for the emergence of bistability: the product of gene *a* inhibits the rate at which *s* is converted into *q*, and its own concentration is itself inhibited by charge *q*. Charge is transmitted back to the electronic device by the biofilm for measurement as electronic current *I*. The process was divided into three main steps: 1) *s* diffuses through biofilm, 2) *s* is consumed and *q* generated by cells, 3) *q* reaches the electrode to generate *I*. **C.** Without the feedback loop (i.e., wildtype cells), the response of *I* to *V* is that of a monostable curve, which increases and asymptotically approaches a maximum current output. **D.** With the addition of the mutually inhibitory synthetic network shown in **B**, a bistable response of *I* to *V* emerges: there is a range of *V* (0.3 in the graph) for which two possible stable steady states of *I* (high and low) are possible.

### Monostable input-output dynamics

The current output (*I*) of an electroactI-Ve biofilm attached to an electrode depends on the potential of the electrode (*V*). By simulating the steady state current output at different potentials, an *I-V* response curve can be obtained like that shown in Figure 1C. This kind of response is typical of an electroactive biofilm of *Geobacter Sulfurreducens* feeding on an a single electron donor substrate, where *I* tends to increase with *V* and asymptotically approaches a maximum current output (*37*). The *I-V* response in Figure 1C was obtained from numerical simulations of the model outlined in Figure 1B. The model includes three important steps in current production. In step one, the electron donor substrate (*s*) diffuses through the biofilm and becomes available to the electrogenic bacteria. Step two includes the oxidation of *s* by the bacterial metabolism, the transfer of an electron *e*^-^ to the electron transport pathway, and the resultant export of charge (*q*) to the surrounding biofilm. In step three, *q* is transported through the biofilm to the electrode, where electrochemical reactions take place at a rate dependent on the electrode potential *V* to generate electrical current *I*. Modeling these three steps can predict the evolution of *I* over time given *V*, and can be used to generate *I-V* response curves as in Figure 1C.

### Bacteria as electro-genetic interfaces

In the model presented here it is the activity of the bacteria that processes the electronic input *V* into the electronic output *I*. Specifically, it is the rate of step two (Figure 1B) that ultimately determines *I* for a given *V*. We can therefore think of the *I-V* response as a biocomputation as shown in Figure 1A. Here electronic inputs and outputs *V* and *I* are transduced into the chemical signal *q* by the biofilm. *q* affects the rate of step two, and the rate at which the bacteria produce *q* to process input into output. In the case of Figure 1C, the transformation of *V* into *I* is relatively simple. However, it is possible that more complicated dependencies of the rate of step two on q, based on synthetic biological networks, could produce a wide variety of different *I-V* responses and be used for more complex biocomputations.

### Bistable input-output dynamics

The electrogenetic toggle switch allows current production *I* of the electroactive biofilm to be switched between high and low states by induction using electrode potential *V*. This switching behaviour is a fundamental step toward more complex computations. To implement the switch the *I-V* response function shown in Figure 1C must be engineered to be bistable as in Figure 1D. In this bistable system, there is a range of *V* for which two possible stable steady states of *I* are possible. For example in Figure 1D, at *V* = 0.3, *I* may be either ‘high’ at around 0.65 or ‘low’ at around 0.05. In order to obtain this bistability we introduced into the model the regulatory dynamics of a gene *a*, whose expression inhibits the rate of step two (i.e., the conversion of substrates to charge q) and whose expression is itself inhibited by high concentrations of charge q. This negative feedback loop admits the emergence of bistability, but not for all values of parameters for the genetic network. That is, not all genetic parts would be suitable for achieving bistability. The range of values of *V* for which two stable steady states exist is the bistable region of the system, and it will be important to determine the existence, size and position of the bistable region for different parameters of the genetic networks in order to engineer a robust switch.

### Mathematical model description

In order to identify parameter sets suitable for the emergence of bistability a mathematical model of the electrogenetic toggle switch was developed. This mathematical model is the reaction-diffusion equation (Equation 1) describing the evolution over time of the concentration of each reactant at each position in the biofilm.

**u**_*x*_ is a state vector tracking the *q, s* and *a* concentrations at position *x* in the biofilm at a given time.

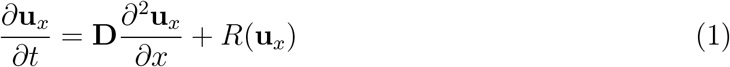

Where **u**_*x*_ is a state vector tracking the *q, s* and *a* concentrations at position *x* in the biofilm at a given time *t*. The individual components of the vector **u**_*x*_ are the state variables as in the following:

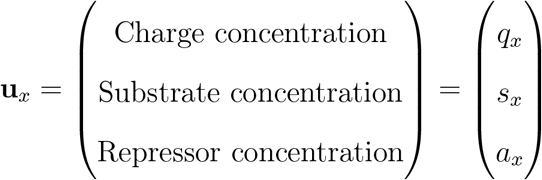

The first term of the right hand side of Equation 1 accounts for transport of the reactants through the biofilm driven by concentration gradients. Where **D** is a diagonal matrix of apparent diffusion coefficients for each of the reactants transport, and reactants tend to move from positions with high concentrations to positions with low concentrations.

The second term *R*(**u**_*x*_) describes how concentrations change due to reactions at each position in the biofilm. In the interior of the biofilm, that is for 0 < *x* < *L*, those are the intracellular reactions related to step two and the genetic network which expresses *a*.

Step two is modeled as occurring in a single step:

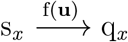

as is the expression and degradation of *a*:

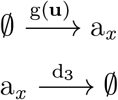

These reactions are described mathematically using the assumption of mass action kinetics in Equation 2.

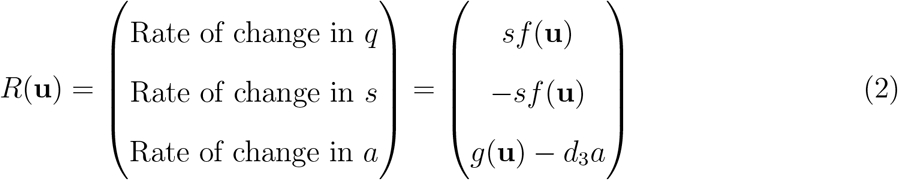

Where the parameter *d*_3_ is the dilution/degradation rate of *a* that balances its expression rate. *f* (**u**) is a hill function describing how the rate of step two changes with the concentration of *a*.

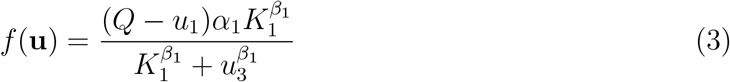

Where *Q* is the maximum capacity of the biofilm for holding charge *q, α*_1_ is the maximum substrate consumption rate, *K*_1_ is the concentration of *a* for which step two is half-maximally inhibited, and *β*_1_ is the hill coefficient of the inhibition. *f* (**u**) tends to zero with higher concentrations of *a*.

*g*(**u**) is another hill function describing how the expression rate of *a* changes with charge concentration *q*. Again, *g*(**u**) tends to zero with higher concentrations of *q*.

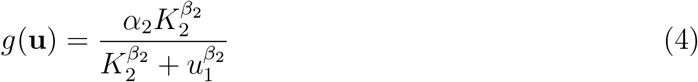

Where *α*_2_ is the maximum expression rate of *a, K*_2_ is the concentration of *q* at which expression of *a* is half-maximally inhibited, and *β*_2_ is the hill coefficient of the inhibition.

At *x* = 0 electrochemical reactions at the electrode-biofilm interface affects the charge concentration, where electrons are exchanged between the biofilm and electrode in both directions at a rate that is proportional to the electrical current at the electrode. This adds an extra term in the expression for the rate of change of *q* in Equation 2.

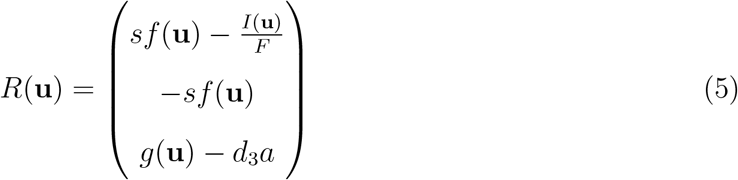

Where *I*(**u**) is the electrical current at the electrode and *F* is the Faraday constant. *I*(**u**) is modeled using the Butler-Volmer relation as outlined in Methods, and depends both on the concentration of charge in the biofilm and potential of the electrode.

### Obtaining bistability in a spatially homogeneous model

Reduction to a spatially homogeneous model reduces dimensionality and makes identification of suitable parameters easier. Spatial homogeneity means that we assume that the state vector **u**_*x*_ is identical for all values of *x*. That is, depth in the biofilm does not influence the concentrations of *q, s* or *a*. If we further assume the concentration of *s* is not limiting, and that *I* is linear in the concentration of *q*, then simplification and nondimensionalisation of Equation 1 yields a pair of coupled ordinary differential equations (ODEs) in two variables.

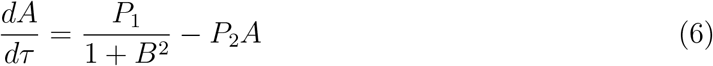

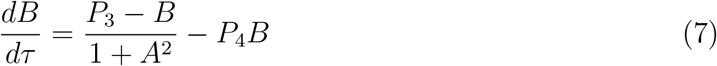

Where the dimensionless variable *A* is a scaled level of expression of *a* and the dimensionless variable *B* is a scaled concentration of charge *q. τ* is dimensionless time, and *P*_1_, *P*_2_, *P*_3_, *P*_4_ are a new set of dimensionless parameters whose values must be selected so as to produce bistability. In this reduced model, desirable parameter values can now more easily be found by inspection of the geometry of the curves shown in Figure 2A. These curves are the nullclines of Equations 6 and 7, which are the points at which the rates of change of *A* and *B* are zero. The intersections of these curves are fixed points of the entire system, and three intersections are required for bistability.

**Figure 2:**
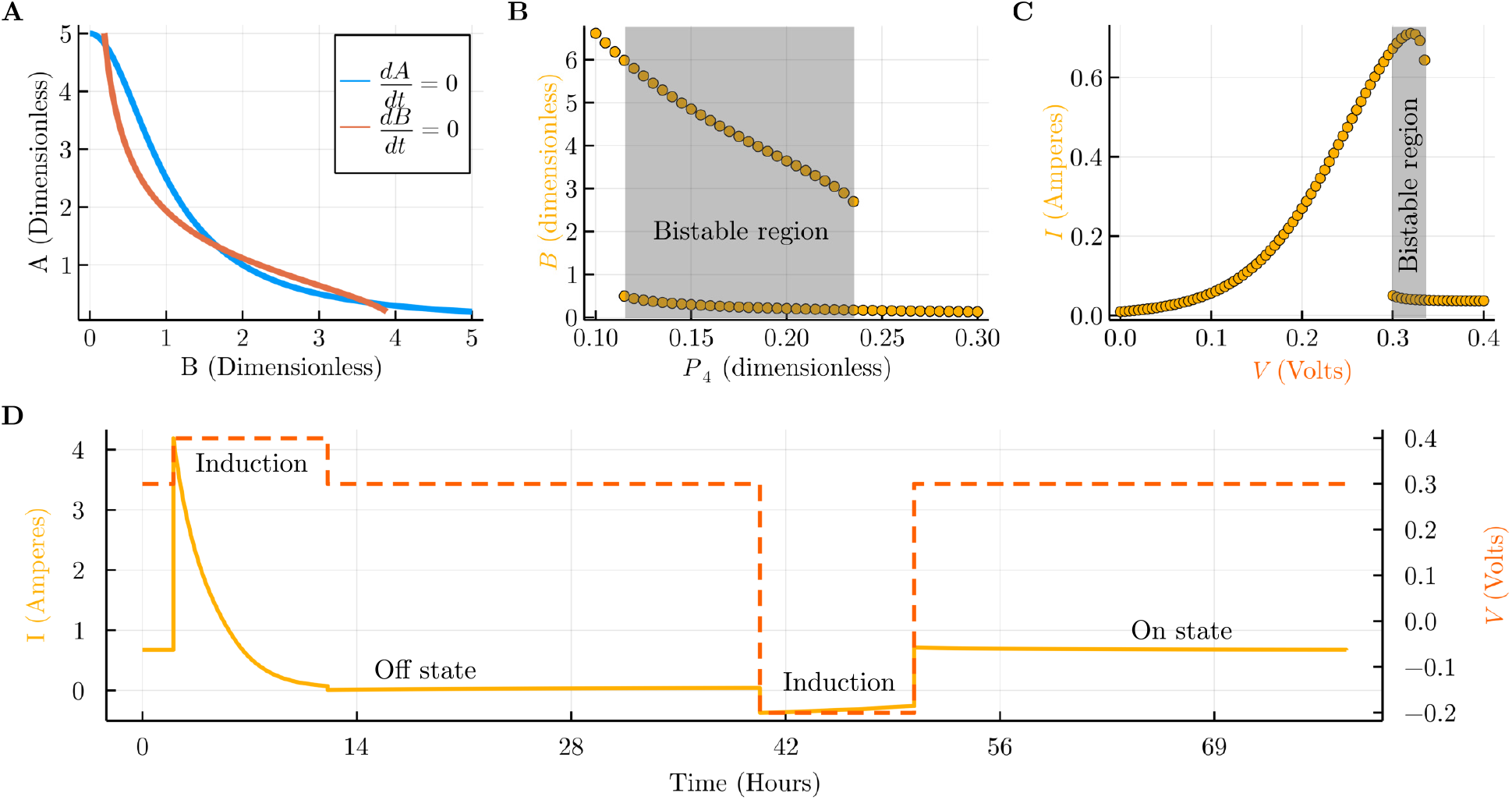
System performance assuming spatial homogeneity. **A.** The nullclines of the dimensionless model of Equations 6 and 7 where the derivatives with respect to each variable are zero. At the intersections of the nullclines are the fixed points (steady states) of the dimensionless model. Bistability requires three intersections, and this can be achieved by inspecting the curves and adjusting the dimensionless parameters. **B.** Bifurcation analysis after parameter adjustment with a bistable region marked in grey. Bistability is shown as the performance of *B* when parameter *P*_4_ changes. **C.** Bistable region in the *I-V* response when these dimensionless parameters were mapped back to the original model parameters. As shown, bistability preserved. **D.** Time course simulation of the model where the electrode potential *V* is used to switch between On / High and Off / Low current states of the switch successfully for this set of parameters.

Fixing the values of the dimensionless parameters such that there are three intersections, as in Figure 2A produces a bistable response in *B* as parameter *P*_4_ is varied (Figure 2B). It also fixes the relationships between the dimensionless parameters that encode design principles which can be followed to obtain bistability in the model with the original parameter set. In particular, in the bistable example from Figure 2A two relationships hold that are related to parameters of the synthetic gene network.

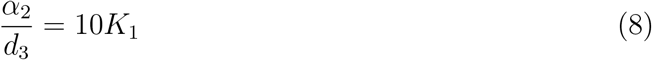

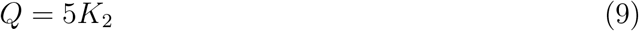

Equation 8 says that *K*_1_, the concentration of *a* that half-maximally inhibits step 2, must be 10 times less than the maximum steady state of *a*. Equation 9 says that *K*_2_, the concentration of *q* that half-maximally inhibits the expression of *a*, should be 5 times less than the maximum capacity of the biofilm for *q*. Using these parameters in the spatially homogeneous model produces a bistability in the *I-V* response curve shown in Figure 2C, whose bistable region starts at around 0.3*V* and ends at around 0.33*V*. The switch then operates as in the time course simulation in Figure 2D. The switch is initially in the ‘On’ state and produces around 0.8*I* at an *V* of 0.3, which is within the bistable region. In order to flip the switch to the ‘Off’ state, *V* is increased to 0.4 for a short time, inducing the system to move into a region of monostability in Figure 2C. After *V* is returned to the bistable region at 0.3, *I* remains low, and the system produces around 0*I* at 0.3*V*. The system will remain in this ‘Off’ state indefinitely, but can be switched back ‘On’ by induction with a temporary step change in V, as is seen at around 40 hours in Figure 2D.

### Challenges due to spatial heterogeneity

Gradients of charge and substrate in the biofilm must be taken into account when engineering the electrogenetic toggle switch (Figure 3). The previous model assumed that the biofilm is a spatially homogeneous environment where gradients of *q* and *s* do not exist. However, previous studies suggest that this assumption is not appropriate (*38, 39*), especially for thicker biofilms of more than around 10*μm* depth. For thicker biofilms, charge transport, substrate diffusion, or both may be limiting the limiting factors for current production *I*. Therefore in practice we can reasonably expect gradients of charge and substrate concentration in the x dimension of the biofilm.

**Figure 3:**
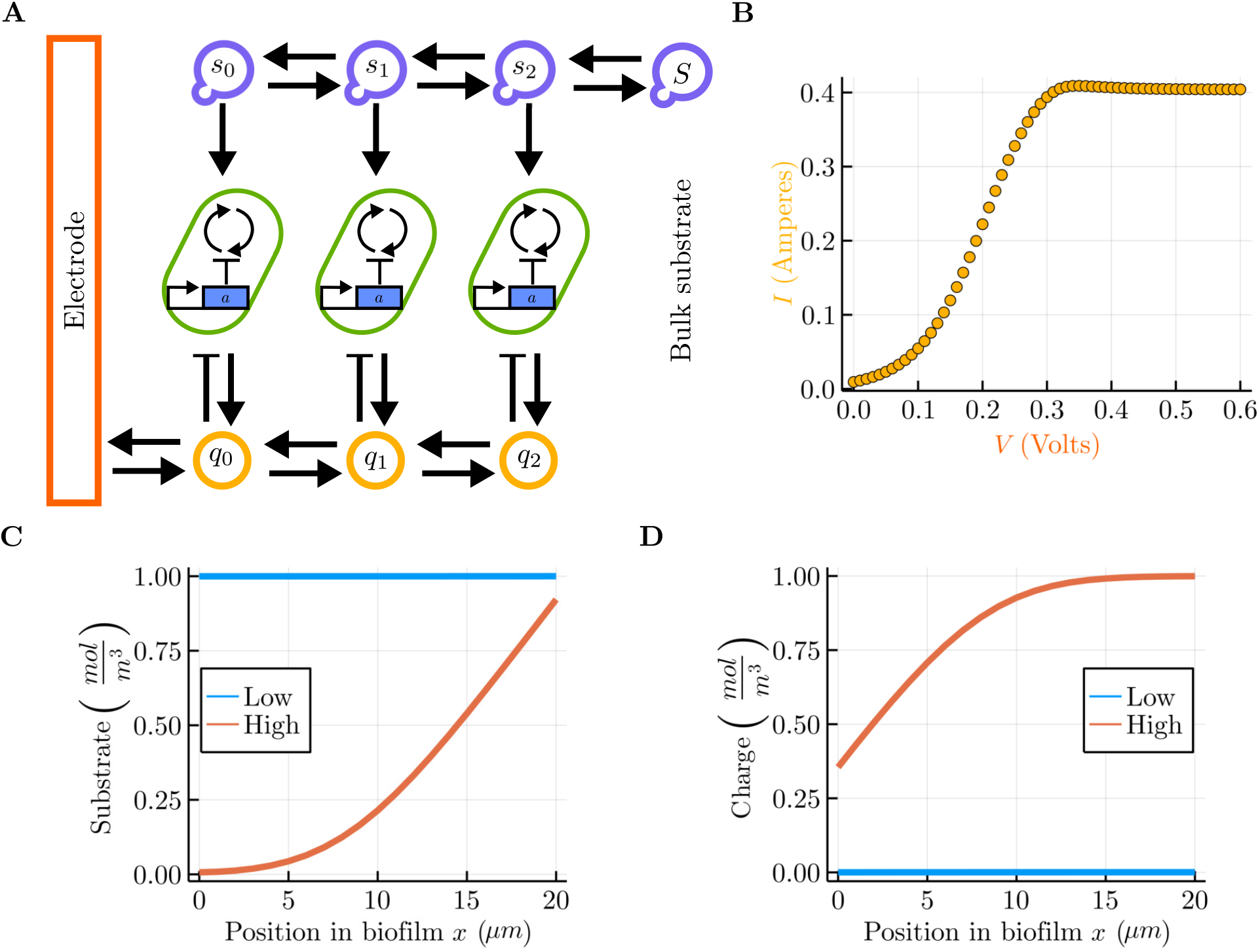
Impact of spatial heterogeneity on switch performance. **A.** Spatial model, where the biofilm was discretised for numerical simulation. Adding a spatial dimension to the homogeneous model incorporates charge (*q*) transport and substrate (*s*) diffusion processes, and allows for the possibility for gradients of charge and substrate to form. **B.** Result of numerical steady state analysis of such a model, showing that the bistability present in the homogenous case can collapse into monostability when spatial gradients are considered. **C.** Dynamics of substrate (*s*) gradients. The blue line plots the gradient obtained in a simulation with low activity of the bacteria (*α*_1_ = 10^-7^) and the orange line plots the gradient with high activity of the bacteria (*α*_1_ = 10^-5^). **D.** Dynamics of charge (*q*) gradients for the same simulations as **C.**.

It is not immediately obvious how this will affect the performance of the switch or whether bistability can emerge in the presence of these gradients. Consideration of biofilm gradients in the electrogenetic toggle switch is complicated by the fact that the magnitude of the gradients are dependent on the parameters of the system. For example, Figure 3C and 3D show how the magnitude of the gradient of *q* and *s* gradient differ depending on the activity of the bacteria (in this case, the parameter α_1_). When this activity is low, there exists almost no gradient of *q* or *s* (blue lines). However, in the case where activity is high, *s* tends to be higher deeper in the biofilm, away from the electrode. The same is true for the gradient of q, although the shape of the gradient differs. Since we expect the electrogenetic toggle switch to have ‘High’ and ‘Low’ activity states, the dependence of the gradient on the activity means that gradients in the toggle switch will be dynamic. That is, gradients will change during the operation of the switch depending on the switch’s state. This added complexity motivates further the development of mathematical models and the numerical analysis of the system as a whole.

In order to identify the impact of biofilm gradients we performed numerical simulations of Equation 1 with the same parameters that produced the bistable dynamics in Figure 2. The result is the monostable *I-V* response shown in Figure 3B and a system which does not operate as a switch. In order to recover bistable dynamics it is necessary to adjust the parameters of the synthetic gene network.

### Obtaining bistability in a spatially heterogeneous model

By adjusting the parameters of the synthetic biological network in the electrogenetic toggle switch, bistable dynamics can be recovered even if gradients exist in the biofilm. We first considered how to adjust parameters for the case of limiting charge transport only. That is, gradients in *q* are possible, but *s* is assumed homogeneous and constant throughout the biofilm. Using the same parameters identified for the homogeneous model, the *I-V* response in the presence of charge gradients is monostable as shown in Figure 4A. We investigated how the parameters of the inhibition of *a* by *q* described by Equation 4 might be changed to recover bistability in this case. Equation 4 has two parameters, the hill-coefficient *β*_2_ and the half-maximal inhibition constant *K*_2_, and is plotted for *β*_2_ = 2 and 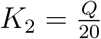 in Figure 4D. With *β_2_ =* 2, we varied K_2_ and plotted the impact on both the shape of Equation 4 and on the *I-V* response of the switch. In Figure 4B and E, *K*_2_ is increased to flatten the inhibition curve in Figure 4E and obtain a bistable region around 0.3*V* — recovering the function of the switch. A further increase in *K*_2_ flattens the inhibition curve further (Figure 4F) and shifts the bistable region to around 0.2*V* but also significantly decreases the size of the bistable region and the robustness of the switch. In these analyses, the bistable region was sensitive to changes in nature of the inhibition between *q* and *a* such that even small changes in the inhibition curve, for example between Figure 4D and E, make the difference between a functional and nonfunctional electrogenetic toggle switch.

**Figure 4:**
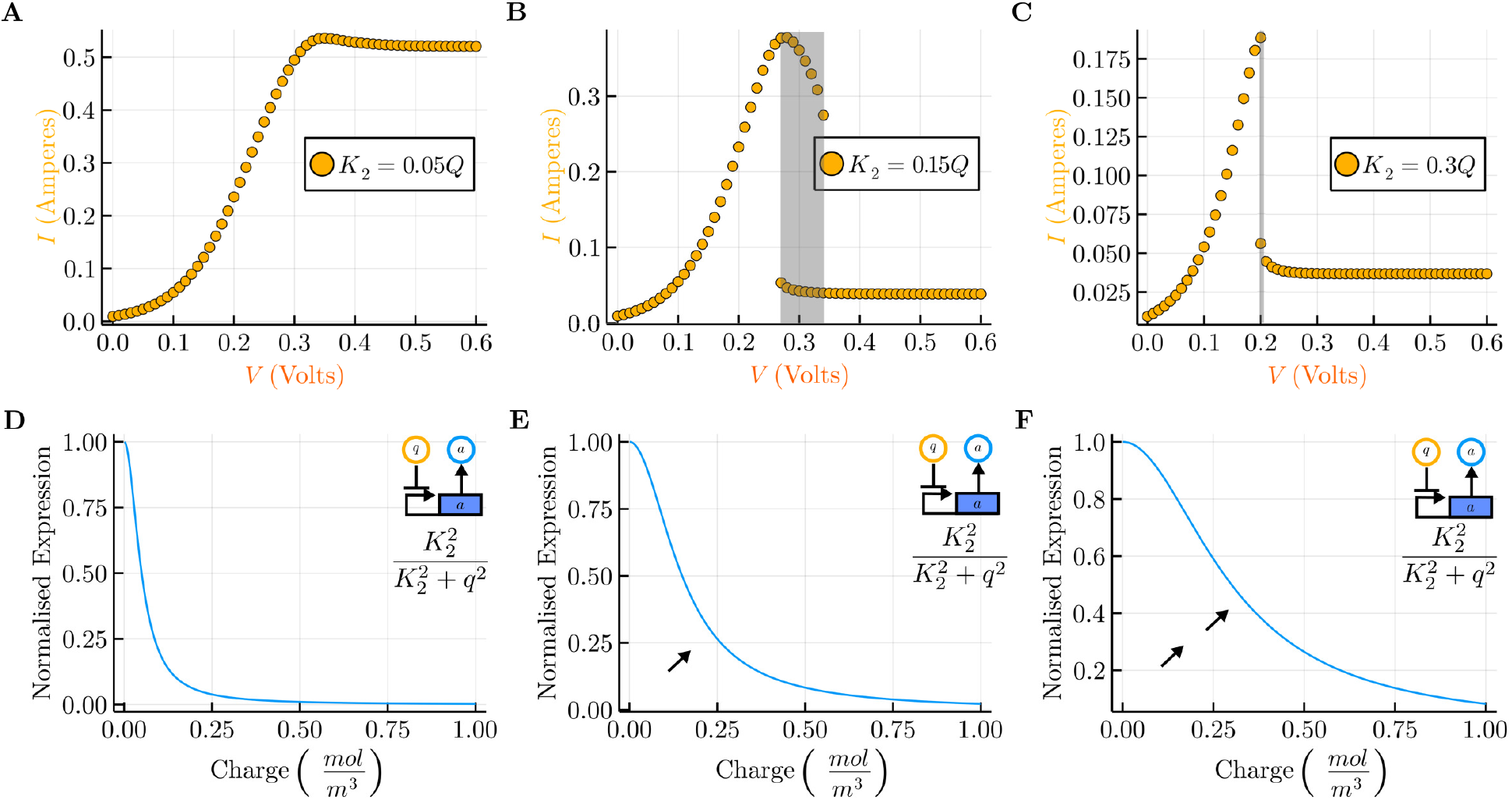
Numerical analysis of the model under limiting charge transport at different values of *K*_2_. The bifurcation diagrams of A, B and C match their corresponding hill-repression transfer functions of plots D, E and F. **A and D.** The bifurcation diagram shows the monostable response obtained when the hill-repression transfer function from *q* to *a* is generated with 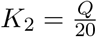. **B and E.** The bifurcation analysis shows how bistability can be recovered by engineering of the genetic circuit such that 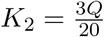, which modifies the transfer function as indicated by the black arrow. **C and F.** The bifurcation diagram shows that the emergence of bistability is sensitive to higher values of the parameter *K*_2_ and the size of the bistable region decreases as the transfer function becomes more linear.

Figure 5 presents a similar numerical analysis for the switch in the case where gradients of both charge *q* and substrate *s* can exist. Figure 5A shows the monostable *I-V* response that is obtained using the parameters from the homogeneous model. As in Figure 4, varying the parameter *K_2_* can recover bistable dynamics. In Figure 5B, there is a qualitatively new kind of behaviour, in that the bistable region extends past reasonable values of *V*. For these parameters, the monostable region that is present at high *V* in Figure 2C disappears. Further increase of *K*_2_ yields bistability with the high *V* monostable region intact, but with a relatively small bistable region.

**Figure 5:**
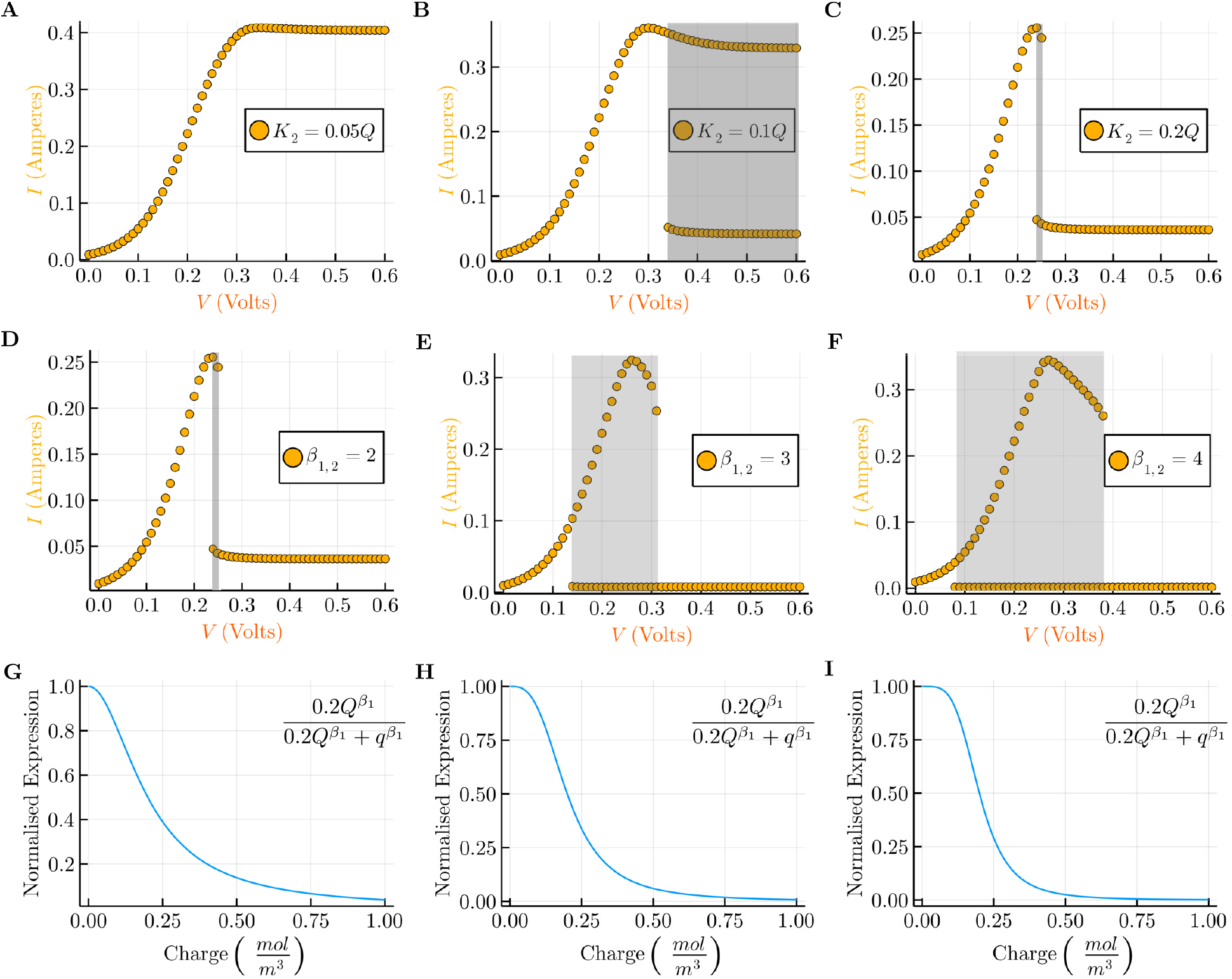
Numerical analysis of the model under both limiting charge and substrate transport in the biofilm. **A, B, C.** The same analysis of *K*_2_ as in Figure 4 but with the addition of limiting substrate diffusion. The result in **B** shows a qualitative change in behaviour, in that for values of *V* over around 0.35 there exists a bistable region that extends to more positive potentials and does not seem to return to monostability for any reasonable value of *V*. **D, E, F** Analysis of the effect of the parameters *β*_1_ and *β*_2_ (the cooperativity of the promoters in the genetic circuit) on the size of the bistable region. Simulations were performed taking the response curve from **C**. The corresponding transfer functions from *q* to *a* are shown in **G, H** and **I**, respectively.

*K*_2_ is not the only parameter of the synthetic biological network that might be engineered. In Figures 5D-F analyses with different values of *β*_1_ and *β*_2_ are shown. These parameters are the hill coefficients of the inhibition interactions between *q* and *a*, and between *a* and the rate of step two. Starting with *β*_1,2_ = 2 and fixing 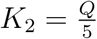 as in Figure 5C, the bistable region grows as *β*_1,2_ is increased in Figures 5E and F. Figure 5E represents the most robust switch found in this analysis, with a large bistable region spanning from around 0. 1*V* to 0.35*V*. This is despite the relatively subtle effect of the parameters *β*_1,2_ on the inhibition interactions, shown in Figures 5G, H and I.

Our analysis predicts that all parameters of the inhibition interactions in the model can be used to tune the performance of the electrogenetic toggle switch. Our initial exploration of this parameter space has predicted that the inhibition interaction characterised by the hill equation plotted in Figure 5I produces the largest bistable region. The higher hill coefficient in this example increases the nonlinearity of the inhibition interaction. Nonlinearity was also an important aspect in the analysis of the original genetic toggle switch, in which the hill coefficient was also an important parameter for increasing the size of the bistable region. Although we found that parameters that admit bistability in a spatially homogeneous model do not necessarily produce bistability in a spatially heterogeneous model, the parameters of the synthetic gene network can be adjusted to recover function.

## 3 Discussion

The model and analyses presented here predict that an electrogenetic toggle switch could be engineered in an electroactive biofilm using a synthetic biological network. The switch produces different steady state current in the ‘High’ and ‘Low’ (or ‘On’ and ‘Off’) states, and can be switched between states using transient changes in electrode potential to which the biofilm is attached. Of fundamental importance to this switch is the feedback loop implemented by the inhibition of gene a by charge *q*, and the inhibition of electrogenic activity by the gene’s expression product (Figure 1B)—the model represents these inhibitions as hill equations. In this case, the electrogenetic toggle switch could be built by designing synthetic gene networks for exoelectrogenic bacteria(*25*) that modify their *I-V* response to be bistable (Figure 1D).

We found it is important to model the impact of spatial heterogeneity that may arise in the biofilm on the performance of the electrogenetic toggle switch. Although heterogeneous conditions are key to the normal function of both single cells(*6*) and biofilms(*40*), spatial homogeneity is an assumption that reduces model complexity and often works well in mathematical models of synthetic biological systems(*41*). However, the electrogenetic toggle switch interacts with the biofilm, and there is good evidence that gradients of charge concentration can exist in electroactive biofilms as a result of charge transport becoming a limiting step in the production of current (*38*). For this reason, we chose to include biofilm transport of substrate and charge in the model, and perform simulations to predict the switch’s performance under conditions where transport is a limiting step. The results presented in Figure 3 show that the same synthetic gene network that produced bistability in the spatially homogeneous biofilm failed to do so in this new context. In the case of limiting charge transport only, and homogeneous distribution of substrate in the biofilm, Figure 4 shows how function can be recovered by adjusting the value of a single parameter in Equation 4. This adjustment represents changes to the genetic components that implement the inhibition of *a* by *q*, using synthetic biological design tools and techniques, that could be guided by the model presented here. For example, the expression of gene *gltA* might be used to control the metabolic rate (current output) of the bacteria (*25*), and the *SoxRS* regulon might be used to sense the electronic signal of the electrode (*23*). In this case, the genetic regulatory components of these genetic networks could be selected, engineered and connected together to provide the mutually inhibitory interactions that are predicted to yield bistability.

The model also predicts that multiple gradients in the biofilm can impact the performance of the electrogenetic toggle switch further. This is especially important since a gradient of one substance in the biofilm can induce a gradient in another. If substrate diffusion through the biofilm is modeled alongside charge transport, we found that performance differs qualitatively (Figures 4B and 5B), despite the same synthetic gene network operating in both contexts. Changes to this network (Figure 5) recover function, and eventually produce a more robust bistable response (Figure 5F). It is also possible that the qualitative behaviour of the switch can change in the presence of multiple gradients in the biofilm. This may cause unwanted behaviours, as in Figure 5B where the *I-V* response is bistable, but only one monostable region exists at electrode potentials below that of the bistable region.

These features—heterogeneity and gradients—build a very dynamic system. The magnitude of the gradients in the biofilm depend on the activity of the electrogenic bacteria, which changes during operation of the switch. Also, the formation of gradients depend on transport in the biofilm being a rate limiting step in current production. However, the magnitude of gradients appear to decrease with the metabolic rate of the electrogens as shown in Figures 3C and 3D, where simulations with high activity of the bacteria produced large biofilm gradients, and simulations with low activity did not exhibit gradients. This coupling of the biofilm gradient to current output has also been observed experimentally (*38*) and needs to be considered for the electrogenetic toggle switch, since we expect the switch to function correctly in both the ‘On’ and ‘Off’ states with high and low electrogenic activity respectively. That is, we expect the switch to operate both in the presence and absence of biofilm gradients. This complexity further motivates the use of modeling and numerical simulation to inform the design of the electrogenetic toggle switch.

Although biofilm depth was a fixed parameter in our model, we carried out initial analysis on the impact of biofilm growth in the performance of the electrogenetic toggle switch (Figure 6). In thicker biofilms we not only expect higher overall levels of electrogenic activity, but also that the charge produced in the deeper parts of the biofilm will take longer to be transported to the electrode. This promotes gradient formation in the biofilm and impacts the performance of the switch as shown in Figure 6, where the size and position of the bistable region changes as the biofilm increases in depth from 10μm to 40μm. The fundamental challenge here is that the biofilm will naturally grow and shrink during operation of the switch, meaning the performance of the switch will change over time. To overcome this methods of controlling biofilm depth could be explored, for example by controlling biofilm formation and dispersion (*42*) using synthetic biological approaches. An alternative is to engineer the bioelectrochemical systems themselves, for example using by using rotating electrodes of flow cells, to control mass transfer rates in the biofilm (*43*). Future work might also consider adding biofilm growth to the model in order to predict which designs are the most robust to changes in the depth of the biofilm.

**Figure 6:**
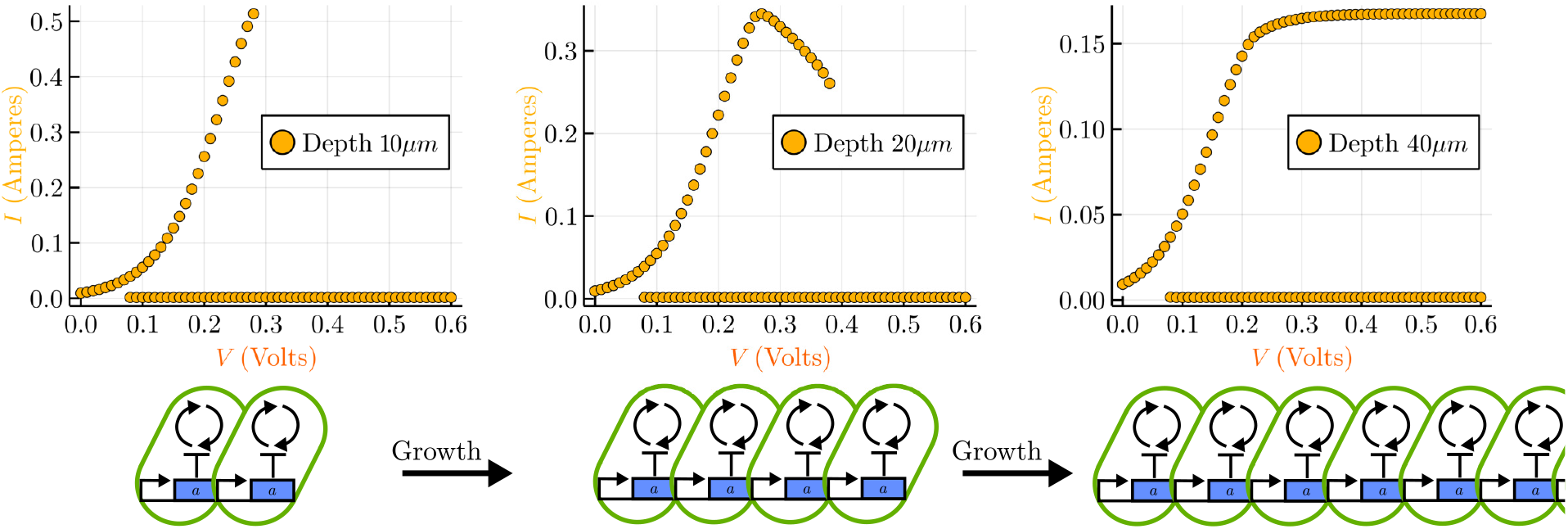
Numerical analysis for the model at various biofilm depths. The simulation of a model of the same genetic circuit and with identical parameters predicts different behaviour depending on the depth of the biofilm. In this example, if the biofilm grow to double in depth from 10*μm* to 20*μm* the size of the bistable region increases. However, doubling again to 40μm produces a qualitative change in behavior, where the system is bistable beyond a certain *V*, but does not return to monostability as for the other biofilm depths. This new behaviour is similar to that shown in Figure 5B.

The model presented here aims to identify general guidelines for engineering an electro-genetic toggle switch. In particular, the focus is on the characteristics of the inhibitions represented in Equations 3 and 4. Previous efforts in building synthetic biological networks to control gene expression with electronic signals (*23*), and vice-versa (*25*), offer a great toolkit for the future implementation of the system. The analysis presented here is a first step toward the rational engineering of a synthetic biological electrogenetic toggle switch.

## 4 Methods

### 4.1 Model formulation

The three-dimensional biofilm was reduced to a single dimension *x*, and is of length *L*. At *x* = 0 is the electrode-biofilm interface and at *x* = L is the interface between the biofilm and the bulk solution. Between *x* = 0 and *x* = *L*, both electron transport and substrate diffusion can occur. It is assumed that the abundance of exoelectrogens and the rates of electron and substrate transport do not depend on *x*.

The diffusion coefficient matrix (*D*) from Equation 1 is a diagonal matrix containing the apparent rates of diffusion for each reactant. Diffusion of *a_x_* is zero for all *x*, since *a* is assumed to be confined to the intracellular environment. Transport of charge and electron holes is balanced, since it is assumed that the concentration of electron holes and charge is conserved, so that for all *x*, 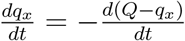 where *Q* is the constant charge capacity of the biofilm.

The partial differential Equation 1 is discretised using the method of lines to produce a system of *N* ordinary differential equations for the numerical analysis. The homogeneous model is simply the case of *N* = 1, for which Equation 1 simplifies to:

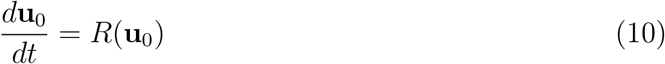

Equations 2 and 5 include three functions of **u**. *f* (**u**) and *g*(**u**) are hill repression functions defined as follows:

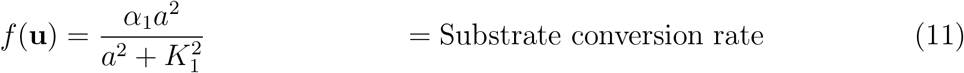

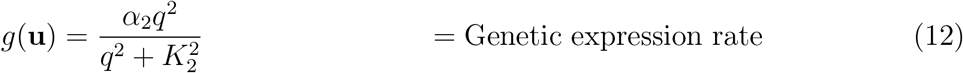

and *I*(**u**) is the current density using the Butler-Volmer relation.

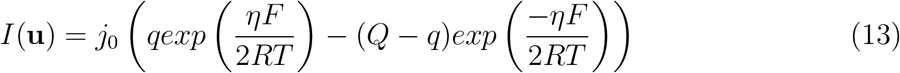

where *j*_0_, *F*, *R, T* and *eta* are the exchange current density, Faraday constant, molar gas constant, temperature, and electrode overpotential.

*R*(**u**) models the coupling of substrate consumption and extracellular electron transport as a single step:

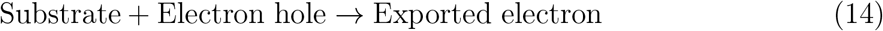

Though in reality it is a result of complex metabolic activity and electron transport machinery (*44*).

The meanings of the parameters introduced in the above equations are summarised in Table 1.

**Table 1:** Parameters of the mathematical model and their default values.

| Parameter | Unit | Value | Description |
| --- | --- | --- | --- |
| $\mathbf{D}$ | $m^2 mol^{-1} s^{-1}$ | - | Diffusion coefficient matrix |
| $D_{1,1}$ | $m^2 mol^{-1} s^{-1}$ | 1 | Diffusion coefficient for charge(45) |
| $D_{2,2}$ | $m^2 mol^{-1} s^{-1}$ | $\frac{5.5}{\Delta x^2}$ | Diffusion coefficient for substrate(45) |
| $D_{3,3}$ | $m^2 mol^{-1} s^{-1}$ | 0 | Diffusion coefficient for repressor |
| $j_0$ | $mol s^{-1}$ | $3 \times 10^{-2}$ | Heterogenous electrochemical rate constant |
| $F$ | $C mol^{-1}$ | 96485.3 | Faraday number |
| $R$ | $J mol^{-1} K^{-1}$ | 8.314 | Molar gas constant |
| $T$ | $K$ | 313 | Temperature |
| $\eta$ | $V$ | - | Overpotential of electrode |
| $Q$ | $mol m^{-3}$ | 10 | Charge capacity of biofilm(33) |
| $\alpha_1$ | $mol m^{-3} s^{-1}$ | - | Maximal substrate conversion rate |
| $\alpha_2$ | $mol m^{-3} s^{-1}$ | - | Maximal genetic expression rate |
| $d_3$ | $s^{-1}$ | - | Degradation/dilution rate of $a$ |
| $K_1$ | $mol m^{-3}$ | - | $a$ concentration at half-maximal repression |
| $K_2$ | $mol m^{-3}$ | - | $q$ concentration at half-maximal repression |
| $\beta_1$ | - | - | Hill coefficient of repression substrate conversion |
| $\beta_2$ | - | - | Hill coefficient of repression of $a$ |

### 4.2 Numerical simulation

For numerical simulation Equation 1 was discretised using the method of lines to obtain a system of ordinary differential equations. A second-order central difference scheme was used to discretise the spatial dimension *x*.

These ordinary differential equations were solved using the Rosenbrock method ‘Rodas5’ implemented in the Julia package ‘OrdinaryDiffEq.jl’.

*I-V* response curves were calculated by finding steady states of the ordinary differential equations for different initial conditions. The steady states themselves were found using a dynamic steady state method, again with the solver ‘Rodas5’.

The code used to generate the figures is made publicly available at ‘https://github.com/Biocomputation-CBGP/An-electrogenetic-toggle-switch-design’, and can be run using Julia 1.7 or greater.

## 5 Acknowledgments

This work was supported by the grants BioSinT-CM (Y2020/TCS-6555) and CONTEXT (Atracción de Talento Program; 2019-T1/BIO-14053) Projects of the Comunidad de Madrid, MULTI-SYSBIO (PID2020-117205GA-I00), and the Severo Ochoa Program for Centres of Excellence in R&D (CEX2020-000999-S) from the Spanish MCIN/AEI /10.13039/5011000 11033.

